# Specific Patterns of Bold Variability Associated with the Processing of Pain Stimuli

**DOI:** 10.1101/157222

**Authors:** Tommaso Costa, Andrea Nani, Jordi Manuello, Ugo Vercelli, Mona-Karina Tatu, Franco Cauda

## Abstract

It is well known that the blood oxygen level dependent (BOLD) signal varies according to task performance and region specificity. This ongoing and fluctuating activity reflects the organization of functional brain networks. Peculiar dynamics of BOLD signal are therefore supposed to characterize brain activity in different conditions. Within this framework, we investigated through a multivoxel pattern analysis whether patterns of BOLD variability convey information that may allow an efficient discrimination between task (i.e., painful stimulation) and rest conditions. We therefore identified the most discriminative brain areas between the two conditions, which turned out to be the anterior insula, dorsal anterior cingulate cortex, posterior insula, the thalamus, and the periaqueductal gray. Then, on the basis of information theory, we calculated the entropy of their different time series. Entropy was found to distribute differently between these brain areas. The posterior insula was found to be is the smaller contributor to the entropy rate, whereas the system formed by the thalamus and periaqueductal gray was found to be the major contributor. Overall, the brain system reaches a higher level of entropy during the rest condition, which suggests that cerebral activity is characterized by a larger informational space when the brain is at rest than when it is engaged in a specific task. Thus, this study provides evidence that: i) the pattern of BOLD variance allow a good discrimination between the conditions of rest and pain stimulation; ii) the discriminative pattern resembles closely that of the functional network that has been called *pain matrix*; iii) brain areas with high and low variability are characterized by a different sample entropy; iv) the entropy rate of cerebral regions can be an insightful parameter to better understand the complex dynamics of the brain.

## Introduction

Averaging-based estimations are statistical techniques commonly used in the field of functional magnetic resonance imaging (fMRI). These procedures try to reduce the variability of the signal under the assumption that this feature reflects mostly noise. However, it has been convincingly shown that BOLD signal variance is task dependent and region specific (Misić, Mills, Taylor, & McIntosh, 2010); therefore, it can carry important information about brain activity (Garrett, Kovacevic, McIntosh, & Grady, 2010, 2013; Garrett, McIntosh, & Grady, 2014). Furthermore, BOLD signal variance appears to increase during brain maturation, a fact that is probably due to a parallel increase in the brain architectural complexity (McIntosh, Kovacevic, & Itier, 2008). Accordingly, it has been found to increase after a recovery from a diffuse cerebral injury, when the brain tries to reorganize itself (Raja Beharelle, Kovačević, McIntosh, & Levine, 2012).

Both computational and empirical works (Garrett et al., 2010; Raja Beharelle et al., 2012) have shown that a system, when it is capable of great power of information processing (so as to elaborate a broad dynamic repertoire), exhibits a wide signal variability (Ponce-Alvarez, He, Hagmann, & Deco, 2015). Indeed, the brain, as it happens in all nonlinear systems, is always in an unstable state “on the edge of criticality” (Gustavo Deco, Jirsa, & McIntosh, 2011; Ghosh, Rho, McIntosh, Kötter, & Jirsa, 2008), changing between many possible states as well as networks configurations. When signal variability is low, the system shows a reduced capacity to switch between several possible states; in contrast, when signal variability is high, the system can easily switch between several different states. Moreover, as we have already said, signal variability is not homogeneously distributed within the brain but exhibits a region-specific and task-related modulation (Misić et al., 2010). In particular, the dynamic modulation required by an active task modifies the “resting state” of the brain, a condition, though, in which the brain is only metaphorically “at rest”.

In fact this ongoing, fluctuating activity reflects the organization of a series of temporally coherent brain networks (G. Deco, Jirsa, McIntosh, Sporns, & Kotter, 2009; Gustavo Deco et al., 2011), which can be active both during rest and during the performance of specific tasks (Laird et al., 2011). These temporally coherent brain networks partially reconfigure themselves during the everyday activity in order to cope with a mutable environment and, consequently, also modify the patterns of BOLD signal variability (Cauda et al., 2012; Katori et al., 2011). Specifically, Katori et al. (2011) have proposed a model according to which the short-term modulation of the synaptic connection modifies the structure of the attractor (i.e., an active state of a cell assembly in a brain network) so as to produce flexible dynamics.

Other fMRI studies have showed that BOLD signal decreases in visual detection tasks and that the magnitude of its variability correlates with the magnitude of the response (He, 2013). Furthermore, interesting results – which have been reported for brain field potentials, membrane potentials and neuronal spiking activity – suggest a reduction of the temporal variance of the BOLD signal during a task compared to the resting condition (He, 2011; Poulet & Petersen, 2008).

Quite recently, a study examined the neural network dynamics of the brain at a large-scale level in order to find out how a task or a stimulus can alter the brain spontaneous activity (Ponce-Alvarez et al., 2015). The model proposed by the authors allows to calculate the statistics of brain activity both in the rest and task conditions with respect to a given set of cerebral regions. The study provided evidence for a decrease of the BOLD signal variance in the task condition. These statistical changes have intriguing consequences, as they imply that brain activity is characterized by a larger informational space when the brain is at rest than when it is engaged in a specific task.

Given these premises we aimed i) to test whether patterns of BOLD variability contain information that may lead to an efficient discrimination between task and rest conditions (so as to identify the brain areas that are more informative); and ii) to provide a model of the BOLD dynamics using the framework of information theory, so as to obtain a better understanding of its nature.

To achieve this second aim, we employed a discrete Markov chain model (Applebaum, 1996), which allowed us to describe the difference between task and rest condition in terms of permutation entropy and of occupation of the different state in the phase space of the system. A Markov chain model is a stochastic process having the property that the future state of the process depends only on the present state of the system. The advantage of this approach is that it is possible to model the dynamics of the system using simple matrix algebra on the transition probabilities of the system. In fact, the matrix of transition probabilities provides a compact and unique description of the behavior of the system under investigation, where each element of the matrix represents the probability of transition from a given state to the next one. The idea at the basis of this phase state approach is well illustrated by Figure 1, which shows the BOLD signal trajectories of different brain areas in different conditions.

**Figure 1.**
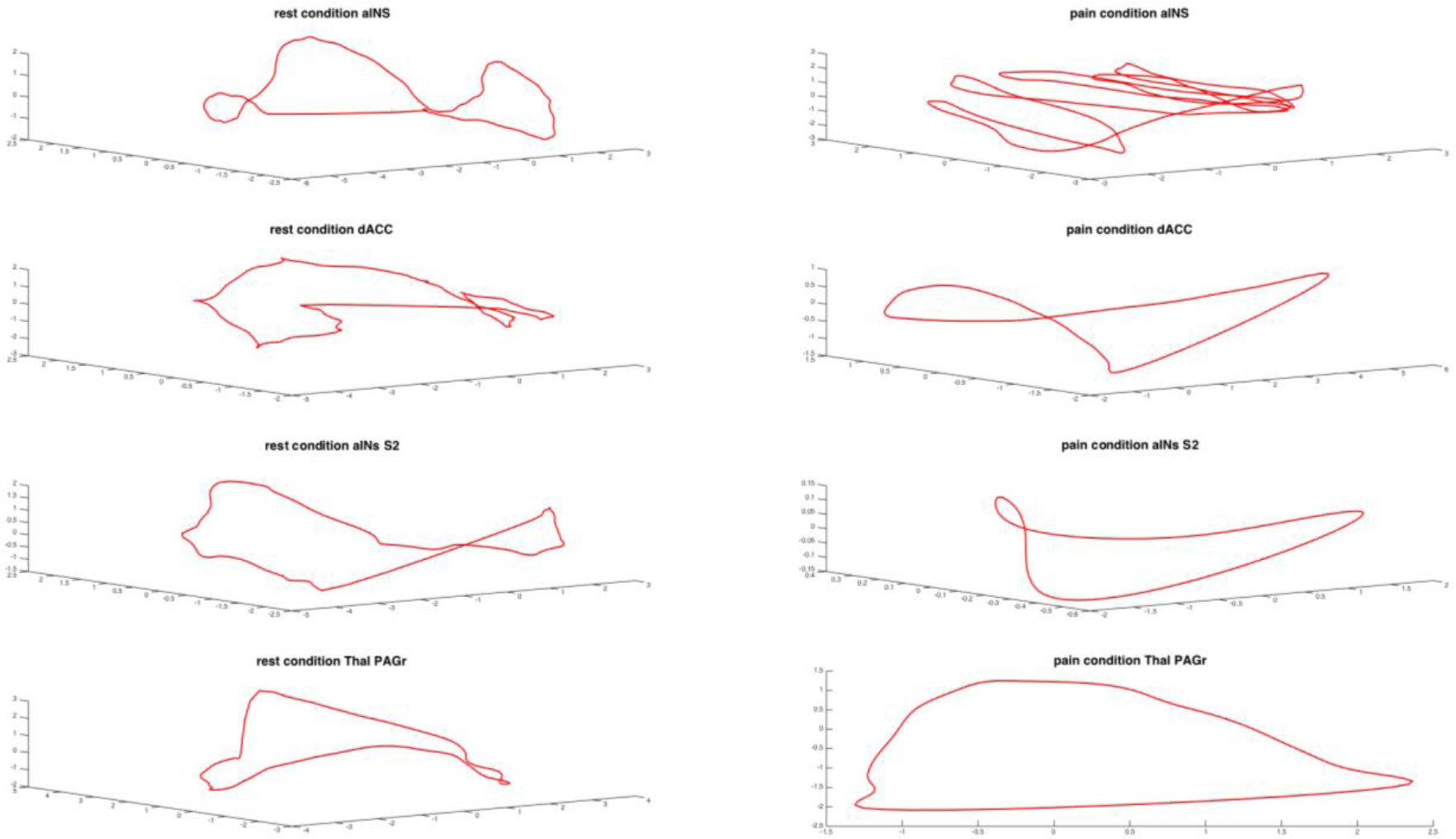
Examples of phase space dynamics for different brain areas in rest and task conditions. During the task condition it is apparent a contraction of the brain dynamics.

## MATERIALS AND METHODS

### SUBJECTS

Seventeen healthy right-handed volunteers (8 females, 9 males, mean age 28±4.2), free of neurological or psychiatric disorders, with no history of drug or alcohol abuse and without medication, participated in the study. Written informed consent was obtained from each subject, in accordance with the Declaration of Helsinki. The study was approved by our institutional Committee of the University of Turin on ethical use of human subjects.

### ACQUISITIONS OF IMAGES

In order to test the dynamics of the BOLD signal, all the 17 subjects were submitted to a mechanical nociceptive task. This procedure has been already used previously by our research group and proved to be a fruitful task condition (Cauda et al., 2014).

Specifically, subjects were engaged in two different experiments.

i) A slow event-related paradigm in which each run was composed of a sequence of 24 brief stimuli of a duration of approximately one second. The interstimulus interval (ISI) ranged pseudo-randomly between 18 and 22 seconds. Four runs of stimuli were applied to the right and left participants’ hands with a hand-held 256 mN pinprick probe stimulator; the location of the stimulation was slightly changed after each stimulus and no more than three consecutive stimuli were applied to the same hand. This kind of mechanical pain is primarily mediated by high-threshold mechanoreceptors and type-I mechano-heat receptors (Magerl, 2001); the pinprick probe is currently used for the evaluation of mechanical pain in both healthy (Baumgärtner et al., 2010) and clinical (Rolke et al., 2006) populations. Images were gathered on a 1.5 Tesla INTERA™ scanner (Philips Medical Systems). All the functional T2* weighted images were acquired using echoplanar (EPI) sequences, with a repetition time (TR) of 2000 ms, an echo time (TE) of 50 ms, and a 90° flip angle. The acquisition matrix was 64 × 64, with a 200 mm field of view (FoV). A total of 240 volumes were acquired, with each volume consisting of 19 axial slices; slice thickness was 4.5 mm with a 0.5 mm gap. Two scans were added at the beginning of functional scanning to reach a steady-state magnetization before acquiring. A set of three-dimensional high-resolution T1-weighted structural images was acquired, using a Fast Field Echo (FFE) sequence, with a 25 ms TR, an ultra-short TE, and a 30° flip angle. The acquisition matrix was 256 × 256, and the FoV was 256 mm. The set consisted of 160 contiguous sagittal images covering the whole brain.
ii) A resting state scan in which the subjects were instructed to keep their eyes closed, while thinking of nothing in particular and not falling asleep. The rest session took 15 minutes with the same parameter of the task session.

### DATA ANALYSIS

The datasets were pre-processed and analyzed using the BrainVoyager QX software (Brain Innovation, Maastricht, The Netherland) and in-house developed MATLAB scripts. Functional images were pre-processed as follows: (i) mean intensity adjustment; (ii) 3D motion correction; (iii) slice scan time correction; (iv) spatial smoothing (FWHM of 8 mm); and (v) temporal filters: linear and non-linear trend removal through a temporal high pass filter (f<3 cycles in time course). After pre-processing, the dataset of each subject was transformed into Talairach space.

#### Multivoxel Pattern Analysis

Specific standard deviation maps of participants were calculated using a MATLAB custom script and then submitted to multivoxel pattern analysis (Tong & Pratte, 2012). Before the train/test step, a reduction procedure employing a recursive feature elimination (RFE) (De Martino et al., 2008) permitted to reduce the number of features to the 6% that retained the highest discrimination power. On the complete set of measured voxels we applied iteratively a training algorithm (least square support vector machine, ls-SVM) to eliminate irrelevant voxels and to estimate the informative spatial patterns. So, the correct classification of the test data increased, while features/voxels were pruned on the basis of their discrimination ability.

For each run, maps were labeled as “rest” or “painful stimulation” on the basis of their membership to rest or stimulation task and then analyzed using the ls-SVM classification algorithm. The two classes of maps were divided into a training and a test set. The training set was used for estimating the maximally discriminative pattern between anterior and posterior clusters with the iterative algorithm; the test set was used to assess the correctness of classification (Bishop, 2006). In order to obtain estimates about the accuracy of the classifier results, we used a *permutation test.* In a permutation test labels are assigned randomly to the example trials, then the classifier is trained on the task with the (wrong) condition labels and finally the generalization performance is tested with a “one-leave-out” cross-validation strategy. Repeating this procedure for 500 times provide a *null distribution* to which the accuracy values, obtained from separate test data, can be compared. Tables were obtained using ClassTal (D’Agata, 2011).

After this analysis, the most discriminative areas turned out to be: anterior insula (aINS); dorsal anterior cingulate cortex (dACC); posterior insula (pINS); the thalami (Thal), and the periaqueductal grey (PAG). From each of these areas we extracted the time series representing the dynamics of the rest and task conditions, which were subsequently used for the entropy analysis.

#### Entropy calculation

The parameter of entropy is one of the most important measures in information theory. In our case, the entropy of a BOLD signal corresponds to the number of bits required to specify a stimulus condition. And the more the bits are, the more complex the stimulus is. Thus, with regard to a time series, the parameter of entropy can provide insightful information about the complexity of its dynamics.

For each brain area we therefore calculated the entropy of the different time series in the two conditions: rest and stimulation. We used the permutation entropy (PE) (Bandt & Pompe, 2002) to investigate different dynamics. The permutation entropy is a robust indicator both of the complexity of the time series and of the dynamics of the signals in the phase space.

Given a time series *x*(*t*) with *t* = l,…,*T*, we created a vector composed of the D subsequent values (D refers to dimension). The values of each vectors were sorted in an ascending order and a permutation pattern *π* was created.

For the sake of clarity, consider the following example of time series *X* = {3,1,4,1,5}. The embedding matrix of vectors with D=3 is:

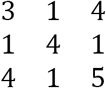

the first vector *s* = (3,1,4) is sorted in ascending order as (1,3,4) and the correspondent permutation pattern is *π* = (102). The second vector is *s* = (1,4,1) and the corresponding permutation pattern is *π* = (021). It must be noted that, if there are equal values in the vector, they are ordered according to their time of appearance. With this method, which can assess the presence or absence of permutation patterns in time series, it is possible to obtain information about the dynamics of the system under observation.

For each permutation *π*, we determined the relative frequency, as follows:

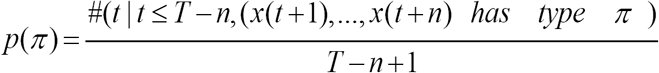

this is an estimation of the frequency of *π*. The PE of order *n* □ 2 is defined as:

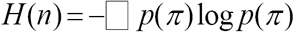

where the sum is over all *n*! permutations *π* of order *n*. So, a completely disordered system is the one that presents an equal probability of having a PE of log*n*!.

To evaluate PE in the two different conditions (i.e., rest and stimulation), we used the relative change formula, given by:

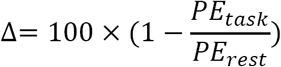

Although entropy is a measure of the response variability, it does not give us any information about whether or not this variability is correlated with changes in the stimulation process or with other unrelated factors. For this reason, we calculated the mutual information, which is the difference between the entropy of the total responses and the entropy of just the task condition. In our case, the entropy of the total responses was given by the sum of the two conditions (rest plus task). This operation removed from the total entropy the contribution not associated with the stimulus (Dayan, 2005).

On the basis of the PE analyses we investigated the dynamics of the time series using the finite Markov chain analysis (Kemeny & Snell, 1960). In a Markov process the outcome at any instant depends exclusively on the preceding outcome, A special kind of Markov process is the Markov chain in which the system can occupy finite or infinite countable states, so that the future of the process depends only on its present state (Markov, 1906). The usefulness of approximating data with a Markov model resides in its flexibility as well as ability to approximate processes that are not truly Markovian.

Employing the same embedding dimension used for the calculation of the PE, we built a phase space for the different time series. In this way we could compare different dynamics. In our case, we used a permutation of order 5 as representation of the phase space.

The dynamics of the phase space was approximated as a finite state of the Markov chain governed by a stochastic matrix *P.* The phase space was divided into a finite number of partitions (*A*_1_,…,*A_n_*) that identified each state of the Markov chain with a subset of the phase space. So, if the phase space is partitioned in n sets, then *P* is a *n* × *n* matrix in which each element (*i,j*) represents the probability that the system goes from state *i* to state *j.*

For each brain area and time series we created a transition matrix of the Markov chain. We then calculated the entropy rate of the Markov chain. Compared to entropy, the entropy rate refers to a measurement of a different statistical feature. In fact, entropy is a property of a random variable, whereas the entropy rate is a property of a stochastic process and quantifies the amount of uncertainty per symbol. The entropy rate is of great interest, because as a measure of an upper bound of information transmission, it provides insight about the type of transmission in different conditions. Furthermore, the entropy rate can measure the degree of complexity of the system taken into consideration. In fact, Pincus (1991) has shown that the approximate entropy of a Markov model is equal to the entropy rate of the Markov chain.

The entropy rate of the associated Markov chain for an ergodic Markov chain, is given by the following formula:

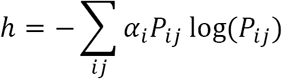

where *α_i_* is the i component of the stationary distribution of the transition matrix *P_ij_* of the process. The entropy rate *h* is the minimal amount of information that is necessary to describe the process.

## RESULTS

The Multivoxel pattern analysis produced a 70% of correct classifications between rest and task condition. The pattern of brain BOLD variability is significantly modified by the painful stimulation; this result allows a good discrimination between the two datasets.

The two patterns of predictive voxels are shown in Fig. 1 and Table 1. Interestingly, the predictive pattern for pain is mostly located in dorsal somatosensory, premotor and sensorimotor areas, whereas the rest of the pattern is located in ventral and secondary somatosensory areas.

The predictive pattern related to rest includes some of the areas (i.e., precuneus and ventromedial prefrontal cortex) that appear to be commonly deactivated in active tasks. There is a evident lateralization of the insular cortex, which is in line to what has been reported in other works of our research group (Cauda et al., 2011, 2012, 2013, 2014; Diano et al., 2016). Of note, only the right anterior insula is involved in the pattern associated with rest.

Taken together, the two patterns comprise some of the areas that are commonly described as parts of the so called “pain matrix” (Friebel, Eickhoff, & Lotze, 2011); however, a GLM analysis on the same data was not able to pick out large portions of these areas (see Fig. 2 lower panels).

**Figure 2.**
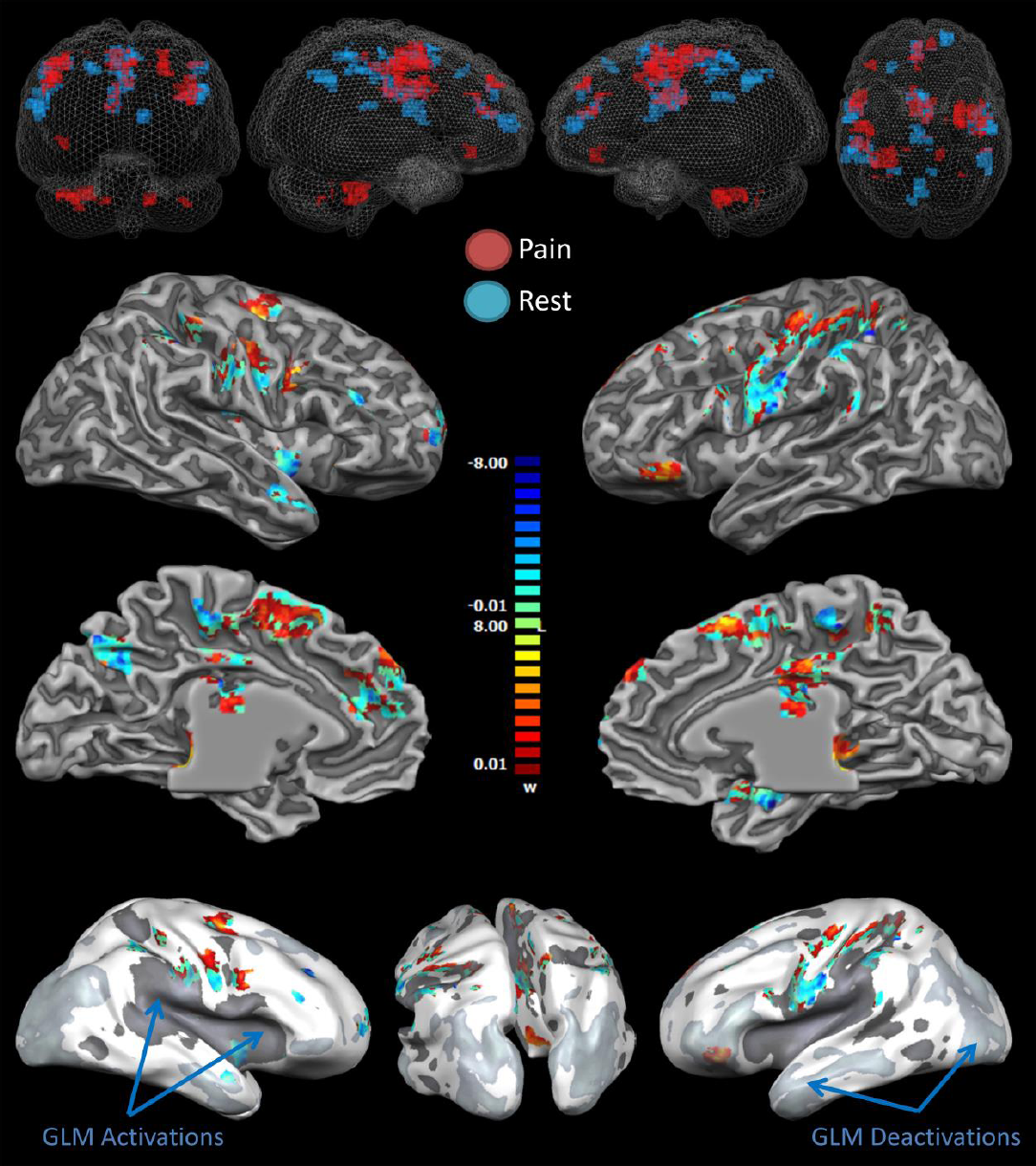
Multivoxel pattern analysis (MVPA) of predictive patterns of variability for pain and rest conditions. The upper panel shows predictive clusters color-coded in relation to the their predictive power: red for “pain” and blue for “rest”. The middle and lower panels show the results of the MVPA. Predictive patterns for pain (colors ranging from red to yellow indicate increasing predictive values) and rest (colors ranging from green to blue indicate increasing predictive values) are showed in 2D and 3D images. Images are visualized with the BrainVoyager QX 3D volume tools.

The PE in the stimulus and resting condition is different for all the investigated areas. Figure 3 shows that the relative change of the PE is higher in the resting condition. This is in agreement with previous works that have reported a reduction of the variability in the stimulus condition. Furthermore, in Figure 3 we can observe the increment of entropy for different brain areas. Intriguingly, the anterior insula exhibits the highest increment, while the posterior insula exhibits the lowest one.

**Figure 3.**
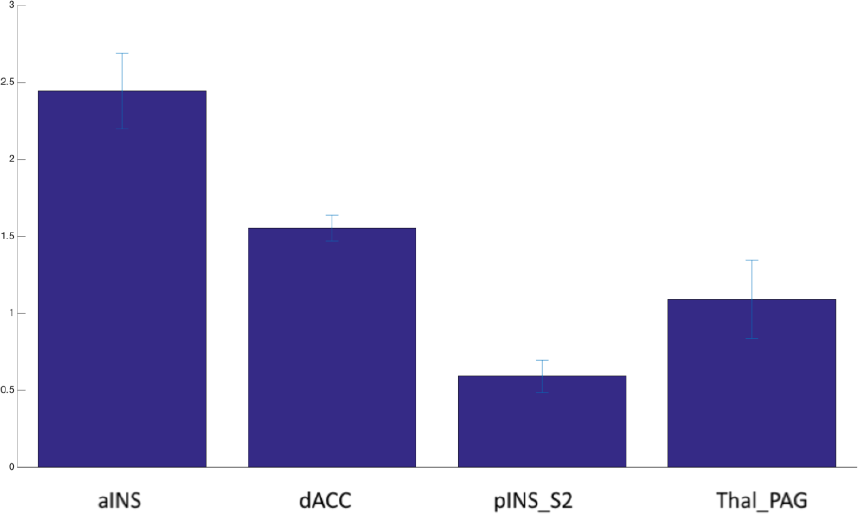
Relative change in permutation entropy. Relative change of permutation entropy between task and rest conditions, in this order. The increment represents an increase of entropy in rest compared to the task condition.

Figure 4 illustrates the relationship between variability and stimulation, which is measured with the mutual information. As shown, the relative change of mutual information in the different brain areas ranges from 6% to 12%. This implies that brain areas are differently activated during the stimulation process.

**Figure 4.**
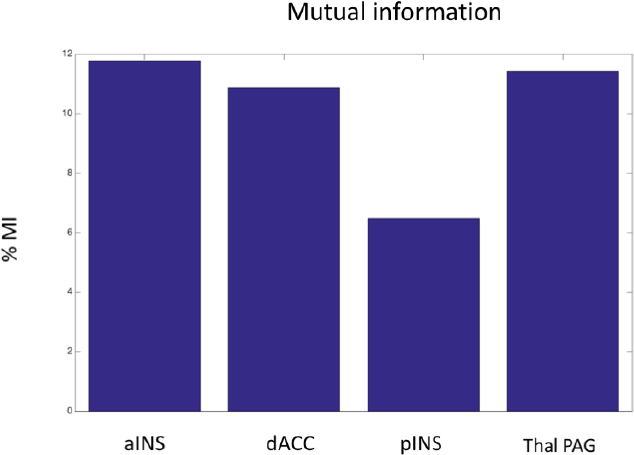
The mutual information for the most discriminative brain areas. Approximately the rate varies between 7% and 12%. The anterior insula exhibits the highest rate, while the posterior insula the lowest.

Figure 5 reports the increment of entropy rate. As we can see, in the resting condition the entropy rate is higher than in the task condition. Different values of increment in the entropy rate range from 0.4 to 0.8, which means that entropy distributes differently between brain areas. The posterior insula is the smaller contributor to the entropy rate, whereas the Thal_PAG system is the major contributor.

**Figure 5.**
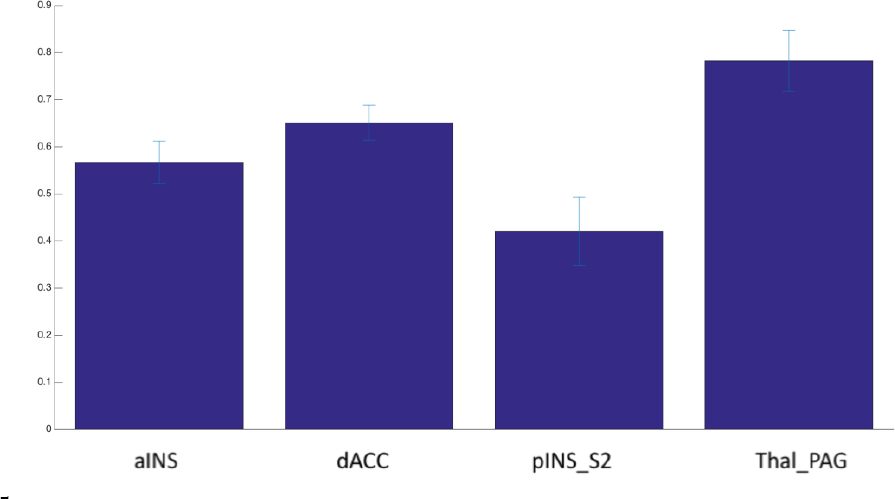
Entropy rate increment for the most discriminative brain areas, which shows that in the resting state condition the entropy rate is higher than in the task condition.

Further information can be obtained from the vector stationary distribution of the Markov chain, a transition matrix and its first time passage for each state of the process. Figure 6 illustrates the transition matrix for our most discriminative brain areas. It is clearly evident the different occupation state between rest and task condition. During the task condition, all the brain areas taken into consideration present a less number of states compared to the resting state condition, which means that in the resting state condition the system covers a greater volume of the state space. This also indicates that, in the resting state condition, the system reaches a higher level of entropy.

**Figure 6.**
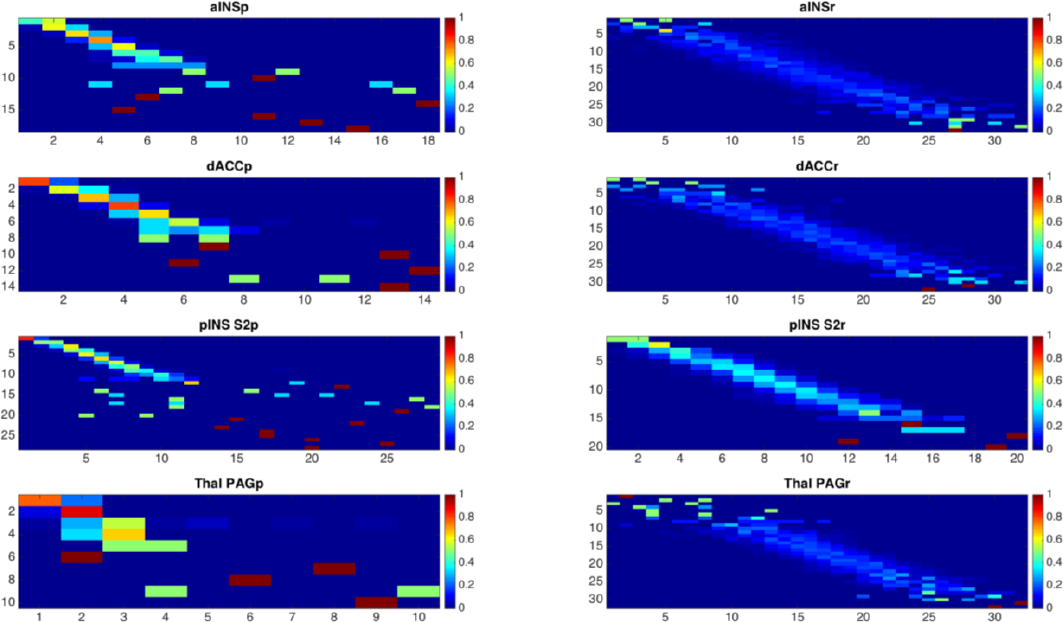
Transition matrix of the most discriminative brain areas in the resting state and task conditions. It is apparent how the phase space is differently covered in the two conditions. During rest, the phase space of the brain system is more extended than in the task condition.

In contrast to the stationary distribution vector, the mean first time passages give us information about the short-range behavior of the chain. Figure 7 shows examples of mean first time passages in both task and rest conditions. In the resting state condition there appear to be more regular and symmetrical transitions between states than in the task condition. This is probably due to the fact that the brain system moves more orderly and smoothly between different states. In fact, in the rest condition the brain system covers all the possible states while it waits for relevant signals coming from the external environment.

**Figure 7.**
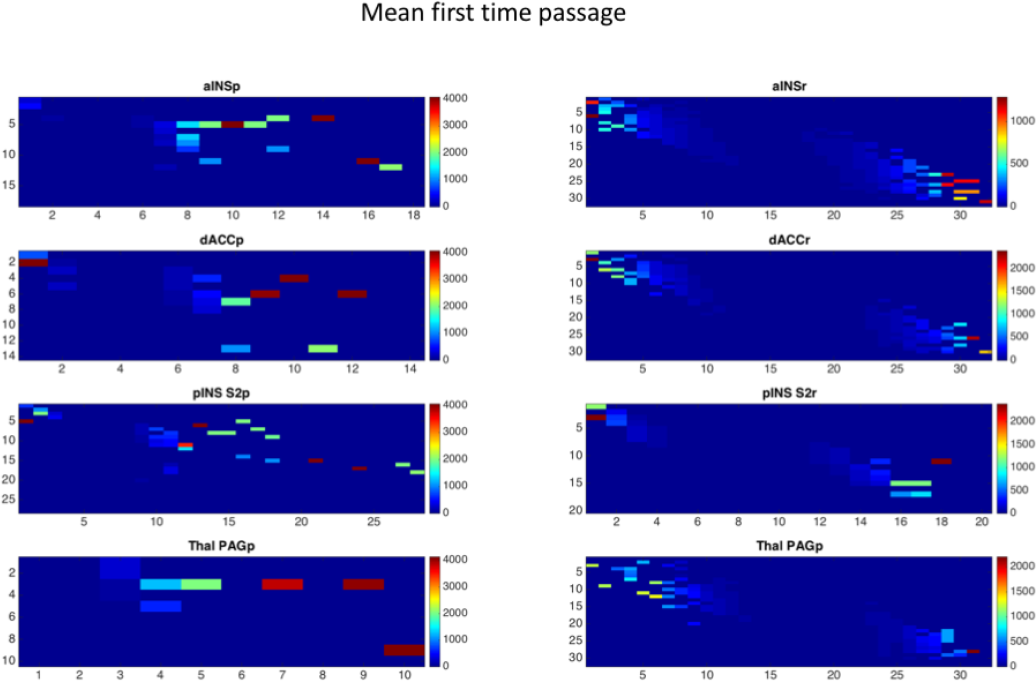
First time passage for rest and task conditions. The mean first passage time from state i to state j is the expected number of steps taken by the process to reach state j from state i for the first time.

Finally, in each condition we calculated the stable limit vector of every transition matrix (see Fig. 8). The stable limit vector represents the limiting distribution of the probability of finding the system in a given state when the system reaches the steady state. The idea of a steady state distribution is that the system has reached (or is converging to) a point in the process where distributions no longer change. At this point forward, all the distributions are equal. So, if the system has reached the limiting distribution and is at a state *i,* the probability that it will pass to an accessible state *j* is the same now and for any time in the future. If they exist, steady states distributions are therefore unique.

**Figure 8.**
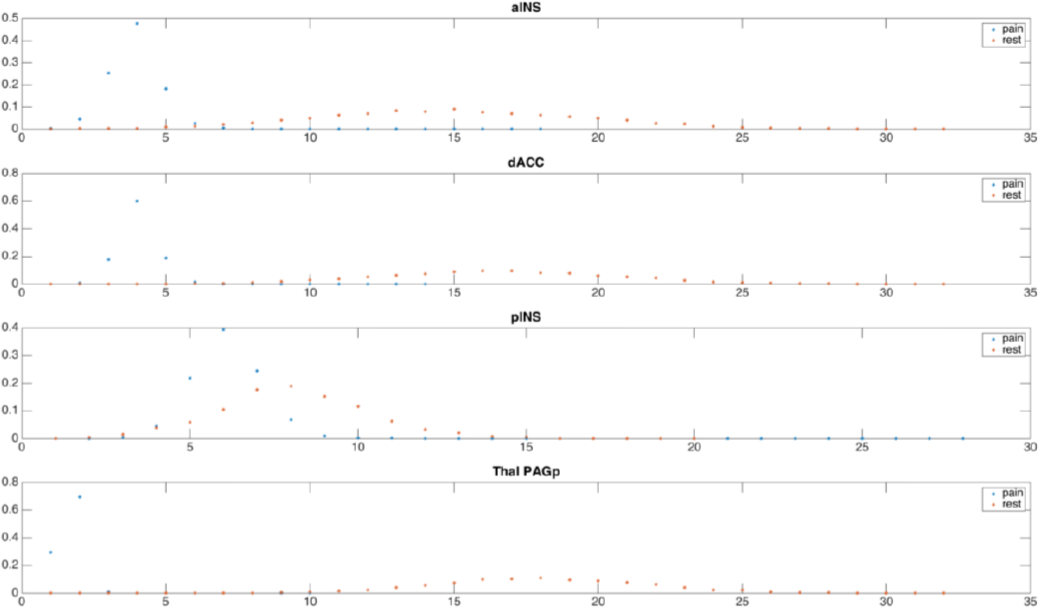
Stable limit vector of each transition matrix for the most discriminative brain areas. For each area, the limit vector behaves differently, both in rest and pain condition.

## DISCUSSION

This study provides evidence that: i) the pattern of BOLD variance permits the discrimination between the conditions of rest and pain stimulation; ii) the discriminative pattern is similar to that of the well known *pain matrix*; iii) brain areas with high and low variability are characterized by a different sample entropy; iv) the phase space of the BOLD signal contains significant information regarding the mesoscopic dynamics of the brain.

The finding that the pattern of BOLD variability can allow us to distinguish between rest and pain perception is consistent with a study by Garrett et al. (2013) about the role and function of the BOLD signal variability that has been able to support the idea that using fMRI it is possible to observe the modulations of this variability in experimental condition-based paradigms. In particular, the authors found different BOLD signal variabilities within and across experimental conditions as well as brain regions, thus showing that signal variability is an effective discriminator between different conditions. The entropy rate appears therefore to be a good parameter in order to distinguish between different brain states, as variability and entropy are proportionally related: high variability is associated with high entropy, and vice versa.

The discriminative pattern between rest and pain perception is also consistent with previous works about the so-called *pain matrix,* as several (though not all, see Fig. 2 lower panels) of the most discriminative areas are part of this functional network. The presence of brain areas that are involved in the pain matrix may be dependent on the methodology, since a painful stimulation can elicit a massive modulation of brain activity, of which only a part is captured by canonical GLM analysis employing predefined BOLD responses.

Overall, brain areas’ entropy, as measured by PE, is lower in the task condition than in the rest condition. This suggests that the brain, when at rest and spontaneously active, generates more entropy as well as more entropy rate than when it is involved in a task. This finding further supports the results obtained by Ponce-Alvarez et al. (2015) and points out that the brain larger activity at rest is likely related to a larger volume of the phase space. In this sense, the observed variability of the BOLD signal is not due to noise but reflects the dynamics of the cerebral system.

The results presented in this study have relevant implications for information processing. As already said in the introduction, several studies have found differences in the temporal variance between young and elderly subjects; this variance is thought to be associated with entropy and condition. The finding that during the task condition we observe a reduction of entropy could be evidence that, in resting state, entropy is an upper bound and is associated with a larger repertoire of states, a repertoire that also contains the state of the task condition and is characterized, therefore, by a greater information capacity, as it is evidenced by its entropy rate. In fact, the entropy rate increases during rest, thus indicating that the state of the brain system has a major information capacity and, as a result, is capable of a greater production and transmission of information.

Within neuroscience (and in particular within the field of neuroimaging) steady-state measurements or simple models of network behavior are commonly assumed to provide a good approximation of the mechanisms underlying sensation, perception, and cognition. However, another view proposes to think of the cerebral cortex as a system in non-equilibrium and of brain computations as unique and transient patterns of activity, elicited by incoming input (Maass, Natschläger, & Markram, 2002). According to this idea, the spatio-temporal structure of the phase space could play a significant role in elaborating information about the external world. Our results, especially the stable limit vector, are in line with this view, as they show that different conditions (rest and task performance) can be clearly distinguished by analyzing the dynamics of brain states.

Differently from Ponce-Alvarez et al. (2015), in this study we focused on the BOLD signals without considering a model of the synaptic activity. But, even if our model is only based on these type of data, its results are consistent with the conclusions of these authors. This provides important evidence that: i) the model of Ponce-Alvarez et al. can be supported by analyses carried out solely on the basis of BOLD signal variability; ii) the entropy rate of cerebral regions can be an insightful parameter to describe mechanisms underling the complex dynamics of the brain.

## CONCLUSIONS

This study has been able to test whether patterns of BOLD variability convey information that may allow to effectively distinguish between task and rest conditions and to provide a model of the BOLD dynamics within the framework of information theory. Specifically, our results show that: i) the pattern of BOLD variance can be a good discriminator between different brain states (task and rest conditions); ii) the most discriminative pattern formed by the areas with high entropy resembles that of the functional network called *pain matrix;* iii) the brain areas with high and low variability are also differently characterized by their sample entropy; and, finally, iv) the phase activity space of the BOLD signal contains valuable information regarding the physiological dynamics of the brain.

**Table 1.**Predictive patterns of variability for pain and rest.

